# Quantifying the Recoverability of V and J Genes from TCR CDR3 Sequences Using Generative Repertoire Models

**DOI:** 10.64898/2026.08.24.746073

**Authors:** Samuel J. Huang, Alexander S. Baras

**Affiliations:** Department of Oncology, Johns Hopkins University School of Medicine, Baltimore, MD, United States; Department of Pathology, Johns Hopkins University School of Medicine, Baltimore, MD, United States

**Keywords:** T cell receptor, immune repertoires, systems immunology, CDR3, V(D)J recombination, generative model

## Abstract

**Introduction:** How much of the variable (V) and joining (J) gene identity of a T-cell receptor is recoverable from its third complementarity-determining region (CDR3) amino-acid sequence alone? Immune repertoire studies often report the CDR3 with V and J annotation that is missing, low-confidence, or inconsistent, so what the CDR3 alone can and cannot fix is both a basic question about the receptor and a practical one for reading those repertoires.

**Methods:** For each of 118,096 pooled human rearrangements (37,687 *α* and 80,409 *β*) we computed the posterior distribution over candidate genes under a generative model of V(D)J recombination and under its post-selection counterpart, and measured recoverability by conditional entropy, the candidate-list size needed to contain the annotated gene, the fraction of sequences admitting a high-confidence single-gene call, and the structure of gene-by-gene confusion.

**Results:** The J gene was nearly determined by the CDR3 in both chains. The V gene was only partially recoverable, and behaved as a group rather than a gene: junctional trimming and non-templated insertion, together with the loss of synonymous codon information in translation, leave sets of mutually confusable V genes whose grouping departs sharply from germline family nomenclature (adjusted Rand index 0.05 for *α* and 0.21 for *β*). Selection sharpened the V posterior modestly (usage-controlled entropy shift −0.06 nats for *α* and −0.28 for *β*) and redistributed which V gene was most probable, a locus-scale rewrite in *β* against a mild reweight in *α*. Both the recoverability measurements and the confusion grouping reproduced in two held-out tumor cohorts.

**Discussion:** V identity is an emergent, system-level property of the repertoire, set jointly by recombination and selection and invisible in any single rearrangement, so it should be reported as a calibrated group rather than a single gene. We also release the pipeline with a computational tool which can output a set of candidate genes with confidence values given a CDR3 sequence.

## 1 Introduction

The diversity of the T-cell receptor (TCR) repertoire arises from V(D)J recombination, where variable (V), diversity (D, in the *β* chain), and joining (J) gene segments are combinatorially joined with random nucleotide deletion and insertion at their junctions [1]. The third complementarity-determining region (CDR3) spans this V(D)J junction and is the most variable part of the receptor and a key determinant of antigen specificity.

The CDR3’s flanking residues are germline-encoded, contributed by the 3*^′^* end of the V gene and the 5*^′^* end of the J gene, while central residues largely arise from the junctional processes [1, 2, 3]. The J contribution is the larger and better characterized of the two, since the J segment contributes a longer and less trimmed germline stretch to the CDR3 than the V segment does. Because the CDR3 is so informative about specificity due to its direct contact with the peptide-major histocompatibility complex, high-throughput repertoire studies routinely sequence and report it, but sometimes without reliable V and J gene annotation, which may be missing, low-confidence, or inconsistent [4]. For example, this may be due to the sequencing read capturing the CDR3 junction but not extending far enough into the flanking germline V and J segments to distinguish among highly similar gene segments [5]. Thus, the amount of V and J identity that can be recovered from amino acid CDR3 is a basic question about the receptor and a practical one for inferring V and J identity when it is absent.

Machine-learning approaches to TCR analysis have grown rapidly, and they largely utilize V and J gene usage as an input feature for specificity prediction or repertoire comparison [6, 7, 8]. Furthermore, some representation-learning methods capture V identity without relying on the CDR3: TCR-VALID, for example, deliberately encodes V identity through the V-specific CDR2 region [9], because the CDR3 alone is insufficient to recover V genes. Recent work has quantified the forward direction of this relationship, showing that V and J gene usage determines on average 80% of CDR3*α* and 65% of CDR3*β* residues [10]. Here, we address the inverse direction, which is how well the V and J genes can be recovered from the CDR3. We present a direct, quantifiable, model-based study of V and J gene identifiability: a posterior distribution over V and J genes given the CDR3, its Shannon entropy, the structure of the residual confusion among genes, and the effect of selection on that posterior.

The junctional processes that erase germline identity act across every gene at once, so the residual ambiguity is a property of the repertoire and only becomes visible when many rearrangements are read together. The question also spans two levels of the same system, because the CDR3s that reach the periphery are produced by the molecular recombination machinery and then shaped by an organism-level selection filter, and the two can be compared directly by reading the same repertoire under a recombination model and under its post-selection counterpart.

A generative model of V(D)J recombination was selected as the tool for this question, because it defines the joint probability of a CDR3 and a candidate gene and yields, by Bayes’ rule, the posterior over V and J genes given the CDR3, which is the exact quantity recoverability requires. We therefore compute this posterior for each CDR3 under OLGA, a generative recombination model [11] built on statistics inferred with IGoR [12], and under its post-selection counterpart obtained by reweighting with the SONNIA selection model [13], so that recoverability can be assessed both before and after selection for the *α* and *β* chains.

The measurements were then repeated in two tumor-infiltrating cohorts that did not contribute to the analysis set: non-small cell lung cancer under anti-PD-1 [14, 15] and head and neck squamous cell carcinoma under anti-PD-L1 and anti-CTLA-4 [16, 17].

## 2 Materials and Methods

### 2.1 Data sources and ingestion

Annotated T-cell receptor sequences were assembled from three public databases: VDJdb [4], NeoTCR [18], and McPAS-TCR [19]. Each dataset was parsed to produce raw TCR records: CDR3 amino-acid string, raw V name, raw J name, and chain. Each record was passed through two ordered normalization steps, and any record failing either step was rejected.

The first step normalized every source gene name to the gene names in the OLGA model [11], so posterior candidates were consistent in terms of naming. For each raw gene name, an ordered list of candidate names in the style of the international ImMunoGeneTics information system (IMGT) was generated from most to least specific: allele suffixes (*01, :01) removed, Adaptive Biotechnologies style (TCRBV. . . ) names [20] rewritten to IMGT style (TRBV. . . ) [2], leading zeros dropped (TRBV01-01 → TRBV1-1), and a family fallback that dropped any trailing “-⟨*n*⟩” subfamily index, and the first candidate present in the OLGA gene set was accepted. Names that matched nothing, and family-only names that were ambiguous against the OLGA set (for example TRAV1 when the model carries TRAV1-1 and TRAV1-2), were also rejected.

The second step normalized the CDR3 to the IMGT junction convention that OLGA expects, namely the segment beginning at the V-region conserved cysteine and ending at the conserved phenylalanine or tryptophan of the F/W-G-X-G motif [3]. The conserved boundary residues were read from the IMGT germline V and J FASTAs [21], with the V cysteine taken as the last cysteine of the V region and the J residue taken from the F/W-G-X-G motif [3]. A CDR3 was retained only if it was non-empty, contained only the twenty standard amino acids, began with cysteine, and ended in the J-germline-appropriate phenylalanine or tryptophan (a generic terminal phenylalanine or tryptophan was accepted when the J-germline residue was unavailable).

Deduplication was performed after normalization, and defined a TCR chain as the full four-tuple of CDR3 amino-acid sequence, V gene, J gene, and chain type. Only TCR chains matching in all four columns would be dropped. After intra-dataset deduplication, the merged TCR *β* dataset consisted of 63,237 chains from VDJdb, 17,666 from McPAS-TCR, and 727 from NeoTCR. Inter-dataset deduplication reduced these to 80,961 unique *β* rearrangements; 665 *β* chains occurred in more than one dataset. The merged TCR *α* dataset consisted of 6,118 chains from McPAS-TCR, 33,081 from VDJdb, and 267 from NeoTCR after intra-dataset deduplication, and inter-dataset deduplication reduced these to 38,971 unique TCR *α* chains; 494 *α* chains occurred in more than one source. At the end of the data processing, 6,134 raw records were rejected (McPAS-*β* 2,429; McPAS-*α* 2,277; NeoTCR-*α* 698; NeoTCR-*β* 229; VDJdb-*α* 410; VDJdb-*β* 91), dominated by gene names that could not be reconciled to the OLGA set (4,936 V and 182 J), CDR3 boundary violations (862), and non-standard residues (154).

### 2.2 Canonical analysis set

The 119,932 unique TCR chains (38,971 *α* and 80,961 *β*) were the input for posterior computation. A further elimination removed sequences whose OLGA generation probability of the CDR3 was zero, which the model treats as impossible and for which both the pre- and post-selection posteriors are empty. This removed 1,284 *α* and 552 *β* sequences, defining a canonical analysis set of *N* = 37,687 (*α*) and *N* = 80,409 (*β*). Every reported Shannon entropy, selection shift, bootstrap, and permutation statistic was computed on this set, defined as the sequences with status “ok” present in both the pre-selection and post-selection arms.

### 2.3 Posterior definition

All posteriors were defined over the OLGA model, taken at gene level after collapsing alleles: 47 V and 61 J genes for the *α* chain and 59 V and 13 J genes for the *β* chain. The *α* model was OLGA’s default human VJ model and the *β* model the default human VDJ model, both built from IGoR-inferred genomic data [12]. Writing *P*_gen_ for the generation probability a recombination model assigns to a rearrangement, the pre-selection posterior was the OLGA prior,

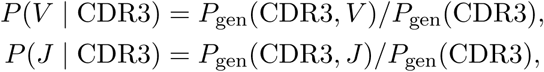

with the opposite gene marginalized by OLGA (computed by masking one gene and leaving the other gene unconstrained); each per-gene weight was the OLGA *P*_gen_ of that masked evaluation, and the weights were normalized within the candidate set so each posterior summed to one. The total *P*_gen_(CDR3) was retained as a flag for impossible sequences. Neither OLGA nor SONNIA receives the annotated V or J gene, which enters only as the evaluation label.

The post-selection posterior multiplied the OLGA generation probability by the selection factor *Q* of the pretrained linear SONNIA model [13]. The pretrained linear *P*_post_ model was used rather than training the deep variant, because fitting a selection model to our own epitope-sorted data would confound the selection factor with the sampling biases of the data under analysis, though the pretrained model carries the sampling characteristics of its own training set.

The linear SONNIA model outputs

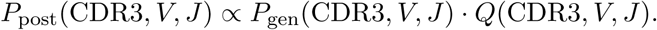

The opposite gene was marginalized over the full *V* × *J* grid,

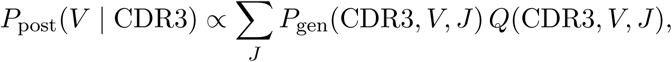

matching the marginalization used in the pre-selection definition. The table was computed for both chains at both gene and family resolution and for the pre- and post-selection arms (Supplementary Table S1). Each posterior was computed once at gene resolution and aggregated to family resolution labels by summing the probabilities of each gene label within a family, which is exact because *P*_gen_ is additive across genes. The family label used the /DV-preserving definition: the locus and family number with the hyphenated member index removed but a /DV⟨*n*⟩ dual-designation suffix retained (TRBV7-2 → TRBV7; TRAV38-2/DV8 → TRAV38/DV8). This single family definition was used wherever a family or group label was required, including the confusion-group analysis below.

### 2.4 Recoverability metrics

Recoverability of a gene from a CDR3 was quantified on the pre-selection posterior with three complementary statistics, each computed at gene resolution and at the /DV-preserving family resolution.

The conditional entropy *H*(axis | CDR3) was the Shannon entropy of a sequence’s posterior in nats, summarized as its mean over the canonical set. Lower entropy denotes a higher confidence in the gene given the CDR3.

Top-*k* recovery was the fraction of sequences whose annotated gene fell within the *k* highest-posterior candidates for *k* from 1 to 20, a measure of the size of the recovered candidate group rather than a measure of pinpoint accuracy of recovery, because many V genes share CDR3 amino acid sequences and cannot be resolved individually. The posterior-mass confidence curve was the fraction of sequences whose top-ranked label carried at least a given posterior mass, swept over thresholds from 0.5 to 0.99 and reported by chain and resolution.

Group structure was examined further by hierarchically clustering the V-by-V posterior-leakage matrix, in which entry (*i, j*) was the mean posterior mass on gene *j* over sequences whose annotated V was gene *i*, restricted to genes with at least twenty sequences. The matrix was made symmetric, converted to distances (one minus similarity), and clustered by average linkage so that mutually confusable genes grouped together. The clustering was then split into the same number of gene groups as there are /DV-preserving V families, so that the data-driven grouping and the family partition contained equal numbers of groups and could be compared on which genes fell together rather than on how many groups each contained. This grouping was compared against the family partition.

### 2.5 Validation framework

The following checks were run:

1. **Independent-source cross-validation.** The *α* analysis was repeated on McPAS-TCR alone, on VDJdb alone, and on the pooled set; NeoTCR-*α*, at 267 chains, was too small to analyze as a standalone source and contributes only to the pooled set. The entropy and grouping estimates agreed across sources that share few rearrangements, and the selection shift kept the same sign and order of magnitude (Supplementary Table S2).
2. **Matched-***n* **subsampling.** The *β* chain was subsampled without replacement to the *α* sample size over 500 replicates and every cross-chain statistic recomputed within each subsample, removing sample size as an explanation for the cross-chain gap (Supplementary Figure S1); the same subsampling underlies the matched-*n* selection comparison reported in Results.
3. **Label-permutation testing.** The pre- and post-selection arms of each sequence were exchanged at random over 2,000 permutations to build a null for the usage-controlled selection shift, which also exposes the estimator’s own finite-sample bias.
4. **Bootstrap intervals.** The canonical *α* set was resampled with replacement over 1,000 replicates, recomputing the entropies, the selection shift and the grouping index on each replicate, and the *β* entropy was bootstrapped at full sample size for comparison.
5. **Held-out calibration.** The candidate-set coverage curve was fitted on half the canonical set and evaluated on the other half, split by a hash of the CDR3; requested and held-out coverage agreed to within 0.4 percentage points for both chains at every level from 0.5 to 0.95.
6. **Held-out cohort replication.** The measurements were recomputed on the two tumor cohorts described in Section 2.6.

### 2.6 Held-out cohort replication

Two single-cell cohorts published after the source databases were used as held-out validation. The first is a 231-patient atlas of anti-PD-1-treated non-small cell lung cancer [14, 15], whose processed data carry paired *α* and *β* annotations for 434,458 cells; the second is a 15-patient head and neck squamous cell carcinoma cohort treated with anti-PD-L1 and anti-CTLA-4 [16, 17], contributing 14,401 *β* clones. Both cohorts are single-cell, while the analysis set is curated bulk and database-deposited receptors.

Every tuple also present in the canonical set was removed. The shared fraction was 3.3% of unique *α* and 0.4% of unique *β* rearrangements for the lung cohort and 0.5% for the head and neck cohort. The remaining rearrangements were subsampled to 10,000 per cohort-chain at a fixed seed to keep the OLGA pass tractable, and the pre-selection V and J posteriors were recomputed from the CDR3 alone under the same models, giving the conditional entropies, top-*k* coverage, confidence fractions and confusion grouping reported in Results. The selection arm was not repeated, because the replication targets the recoverability ceiling and the SONNIA selection factor is a fixed pretrained model that would carry the same training composition into either repertoire.

### 2.7 Statistical tests

The effect of selection on V recoverability was measured as a usage-controlled pre-to-post shift in V-gene conditional entropy. To prevent a mere change in marginal V usage from seeming like sharpening, each post-selection sequence was importance-weighted so that the joint distribution of CDR3 length and post-selection argmax-V matched the pre-selection distribution of CDR3 length and pre-selection argmax-V. Binning sequences by their (CDR3 length, argmax-V) pair, the weight of each such bin was the ratio of its pre-selection to post-selection frequency, and bins never occurring in the pre-selection marginal received zero weight and were dropped. The reported shift was the weighted-mean post-selection entropy minus the mean pre-selection entropy. To confirm the result did not depend on dropping those sequences, the estimate was repeated with an additively smoothed pre-selection distribution, which assigns them a small nonzero weight and retains them.

Generalizing this view from two genes to the whole locus, the redistribution of V identity under selection was characterized for both chains by comparing, per V gene, the pre- and post-selection top-1 share (the fraction of canonical sequences for which the gene was the highest ranked probability) and the marginal usage (the mean posterior mass on the gene), together with the bin-level accounting of the usage control. A gene was termed eliminated as argmax if it was the top-1 call for one or more sequences pre-selection but for none post-selection, and promoted from zero if the reverse.

Sample size was removed as an explanation for the cross-chain gap by subsampling the *β* chain without replacement to the *α* sample size of 37,687 over 500 replicates and recomputing the statistic within each subsample.

Within-chain significance of the *α* and *β* usage-controlled shift was assessed by a label-permutation test where each sequence’s pre- and post-selection arms were independently swapped with probability one half and the usage-controlled shift recomputed, over 2,000 permutations. The two-sided *p*-value was the fraction of permutations whose shift was at least as extreme in absolute value as the observed shift, floored at 1*/*(*n*_perm_ + 1), and the separation was reported in standard deviations of the permutation null.

Confidence intervals were obtained for both chains. The *α* intervals came from resampling the 37,687 canonical *α* sequences with replacement over 1,000 replicates, recomputing on each replicate the three conditional entropies, the usage-controlled shift, and the adjusted Rand index of the data-driven V confusion grouping against the full-data grouping. The usage-controlled shift carries a small finite-sample downward bias whose magnitude is the center of the permutation null, so a bias correction was applied. The bias correction was computed per set from the analyzed set’s own permutation-null center (−0.00147 for the pooled *α* set, not a fixed constant) and applied as a single offset to the bootstrap point estimate and interval, not recomputed within each resample. Because the test unit was the deduplicated rearrangement and residual clonal structure among related sequences could reduce the effective sample size, the permutation *p*-values were treated as lower bounds.

Where a held-out cohort differed from the analysis set in measured recoverability, importance reweighting was used to isolate the contribution of V usage. Each analyzed sequence was weighted by the frequency of its annotated V gene in the cohort divided by the frequency of the same gene in the analysis set. Genes the cohort does not use receive zero weight. Intervals came from resampling the analysis set with replacement over 1,000 replicates at fixed weights.

## 3 Results

### 3.1 The CDR3 determines J but not V, and ***β*** entropy exceeds ***α***

The two genes differed greatly in how completely the CDR3 fixed them. The J gene was nearly determined in both chains, with a mean pre-selection J-gene conditional entropy of 0.0115 nats for *α* and 0.0474 nats for *β*, close to the zero-entropy limit of a fixed gene (Figure 1). The V gene was far less determined and differed markedly between chains. The mean V-gene conditional entropy was 1.2925 nats for *α* against 2.7055 nats for *β*, so the *α* V gene was substantially more recoverable than its *β* counterpart (Figure 1). The *α* advantage held across the full range of CDR3 lengths and was not a consequence of the larger *β* sample. At matched sample size, the *α* entropy of 1.2925 nats remained far below the *β* matched-*n* interval of [2.6986, 2.7127] nats (Supplementary Figure S1), meaning the gap arises from chain biology rather than a sampling artifact. Within the *α* chain, the V-entropy and grouping estimates were similar across the major source datasets and the selection shift kept the same sign and order of magnitude, as summarized in Supplementary Table S2.

**Figure 1:**
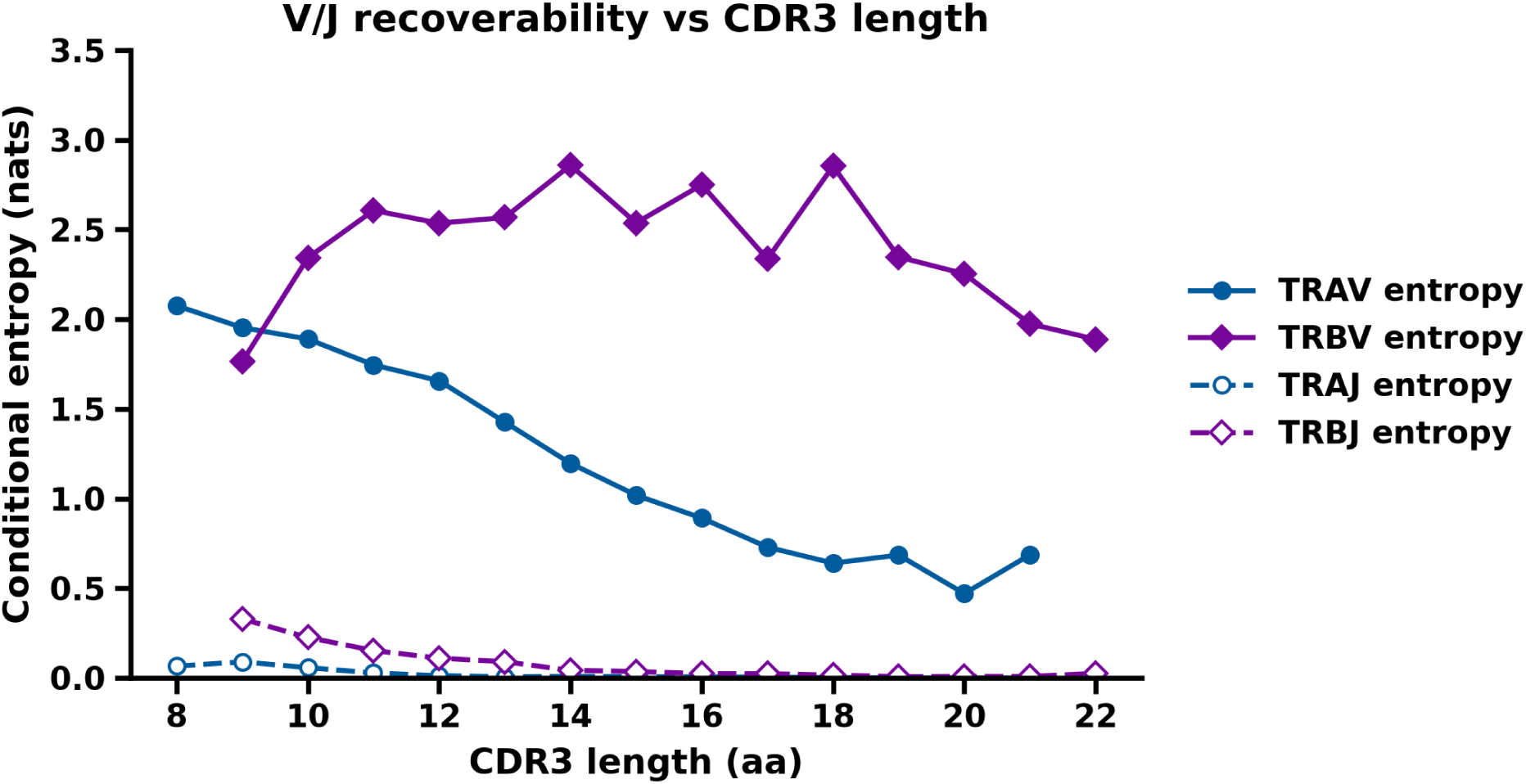
Recoverability of the V and J gene from the CDR3, by chain. Mean posterior conditional entropy (nats) versus CDR3 length for the V gene (upper curves, solid, filled markers) and J gene (lower curves, dashed, open markers), with TRA in blue and TRB in purple as in every other line figure here, on the canonical pooled set (TRA N=37,687, TRB N=80,409; length bins with *n* ≥ 20). The J gene is nearly determined in both chains (mean 0.0115 nats TRA, 0.0474 TRB); the V gene is only partially recoverable and markedly more so for TRA (1.2925 nats) than TRB (2.7055 nats).

### 3.2 Top-***k*** recovery measures candidate-set size, and ***α*** is easier to recover

Each point of the top-*k* curve gave the fraction of sequences whose annotated V fell among the *k* highest-posterior genes (Figure 2). For the *α* chain a single best candidate (Top 1) already contained the true V for 49% of sequences and for Top 10 for 85%, whereas for the *β* chain Top 1 sufficed for only 20% and Top 10 was required to reach 61%. The *α* CDR3 thus ascertained its V gene with a shorter candidate list while the *β* CDR3 needed a substantially longer one to reach comparable coverage, and *α* lay above *β* at every value of *k*. The two curves converged only at large *k*, where both approached saturation (0.92 for *α* and 0.88 for *β* at *k* = 20) as the candidate sets spanned most available genes. This convergence reflected the larger and more diffuse *β* candidate set, not any change in recoverability, since *β* did not overtake *α* at any *k*.

**Figure 2:**
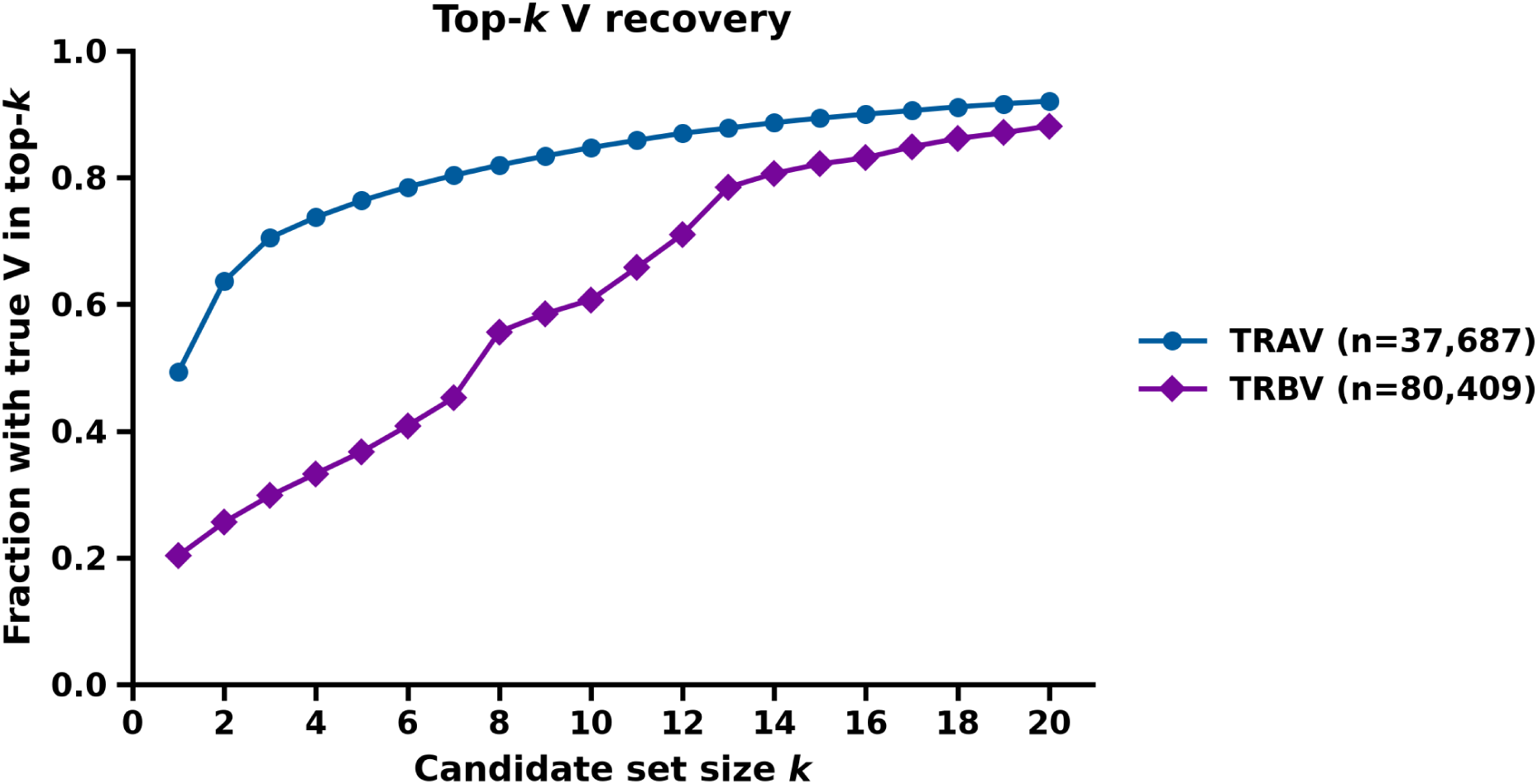
Top-*k* candidate-set coverage for the V gene. Fraction of sequences whose annotated V falls within the *k* highest-posterior genes, k=1..20, TRA vs TRB. A single best candidate contains the true V for 49% of TRA vs 20% of TRB sequences; ten candidates reach 85% (TRA) vs 61% (TRB). TRA lies above TRB at every *k*; the curves converge only near saturation (0.92 vs 0.88 at k=20).

### 3.3 Confusable V gene groups depart from IMGT family nomenclature

The structure governing V-gene recovery did not coincide with the IMGT family nomenclature. Clustering the V-by-V posterior-leakage matrix and forming the same number of gene groups as there are IMGT families produced a data-driven grouping that departed sharply from the family partition, with an adjusted Rand index of 0.0519 for the *α* chain and 0.214 for the *β* chain (Figures 3, 4). The confusable blocks lying on the diagonal of the confusion-ordered matrices crossed family boundaries rather than respecting them in both chains; the near-zero *α* index indicated essentially no correspondence between the two partitions, while *β*’s index of 0.214, though larger, was still low, so *β* confusion only partly tracked the family structure and *α* showed essentially none. The natural unit at which a V gene could be recovered from a CDR3 was therefore a data-driven group of mutually confusable genes and not the IMGT gene family.

**Figure 3:**
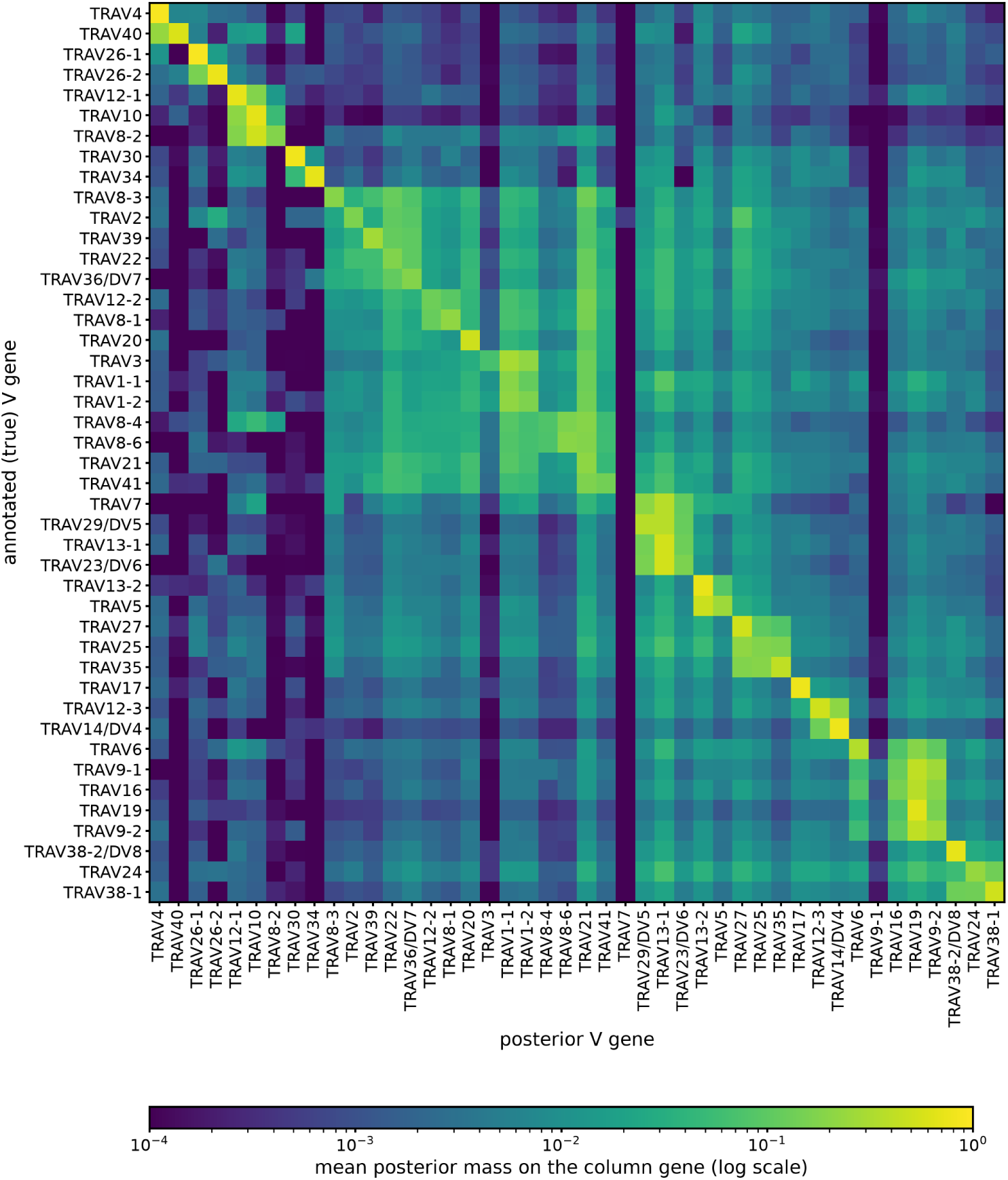
V-by-V posterior confusion matrix, TRA. Rows are the true V gene and columns are the posterior V gene, with each cell showing the mean posterior mass placed on the column gene across sequences whose annotated V is the row gene. Genes with at least twenty sequences are included and are ordered by hierarchical clustering of the confusion structure rather than by family, which places mutually confusable genes adjacent to one another. The bright diagonal blocks are groups of genes that leak posterior mass into one another, and these blocks span multiple IMGT families, so the data-driven grouping departs sharply from the family partition with an adjusted Rand index of 0.0519. Many genes along the diagonal are brightly lit, indicating that a substantial number of TRA V genes retain high posterior mass on the correct gene and are individually or near-individually recoverable.

**Figure 4:**
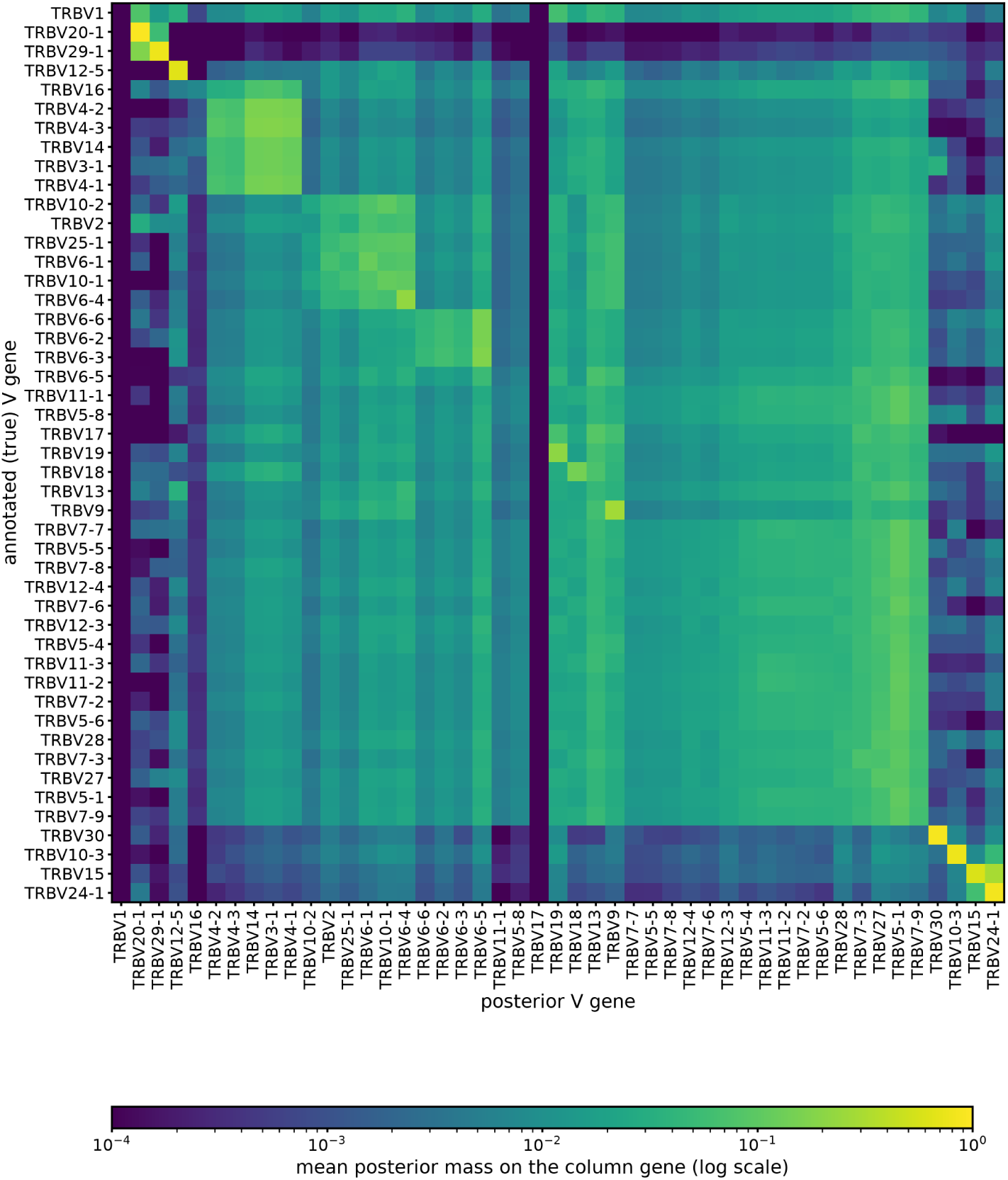
V-by-V posterior confusion matrix, TRB, constructed as in Figure 3 for the beta chain. The confusion blocks only partly track the IMGT families, with an adjusted Rand index of 0.214, so the recoverable unit remains a data-driven group of mutually confusable genes rather than the nomenclatural family. In contrast to the alpha chain, the diagonal is sparse and only a few genes retain strong posterior mass on the correct gene, while most of the matrix is a single broad region of diffuse leakage in which posterior mass is spread widely across many genes. This visual sparsity of the beta diagonal relative to alpha is the qualitative counterpart of the higher beta V-gene conditional entropy and its lower top-*k* recovery.

### 3.4 Confident single-gene V calls are attainable for a majority of ***α*** but a minority of ***β***

A confident single-gene V call was available for a majority of *α* sequences but only a minority of *β* sequences, and for the *β* chain such calls were rare at any usable threshold (Figure 5). Requiring at least half of the posterior mass on the top-ranked V gene qualified 57% of *α* sequences at gene resolution, with 13% still meeting a threshold of 0.99; separately, Supplementary Figure S2 highlights the 18 TRAV genes with mean posterior self-mass ≥ 0.5. The *β* chain was far lower, with only 13% of sequences meeting the 0.5 criterion and 4% meeting 0.99; Supplementary Figure S3 separately shows that 7 TRBV genes had mean posterior self-mass ≥ 0.5. The individual V genes clearing this criterion, meaning those that retain at least half their posterior mass on the correct gene, are listed for both chains in Supplementary Table S3. Coarsening to the family label did not rescue *β* confidence. The *β* family curve sat only marginally above the *β* gene curve at low thresholds (19% versus 13% at 0.5) and coincided with it at strict thresholds. This is consistent with the grouping result that the family is not the unit along which posterior mass concentrates.

**Figure 5:**
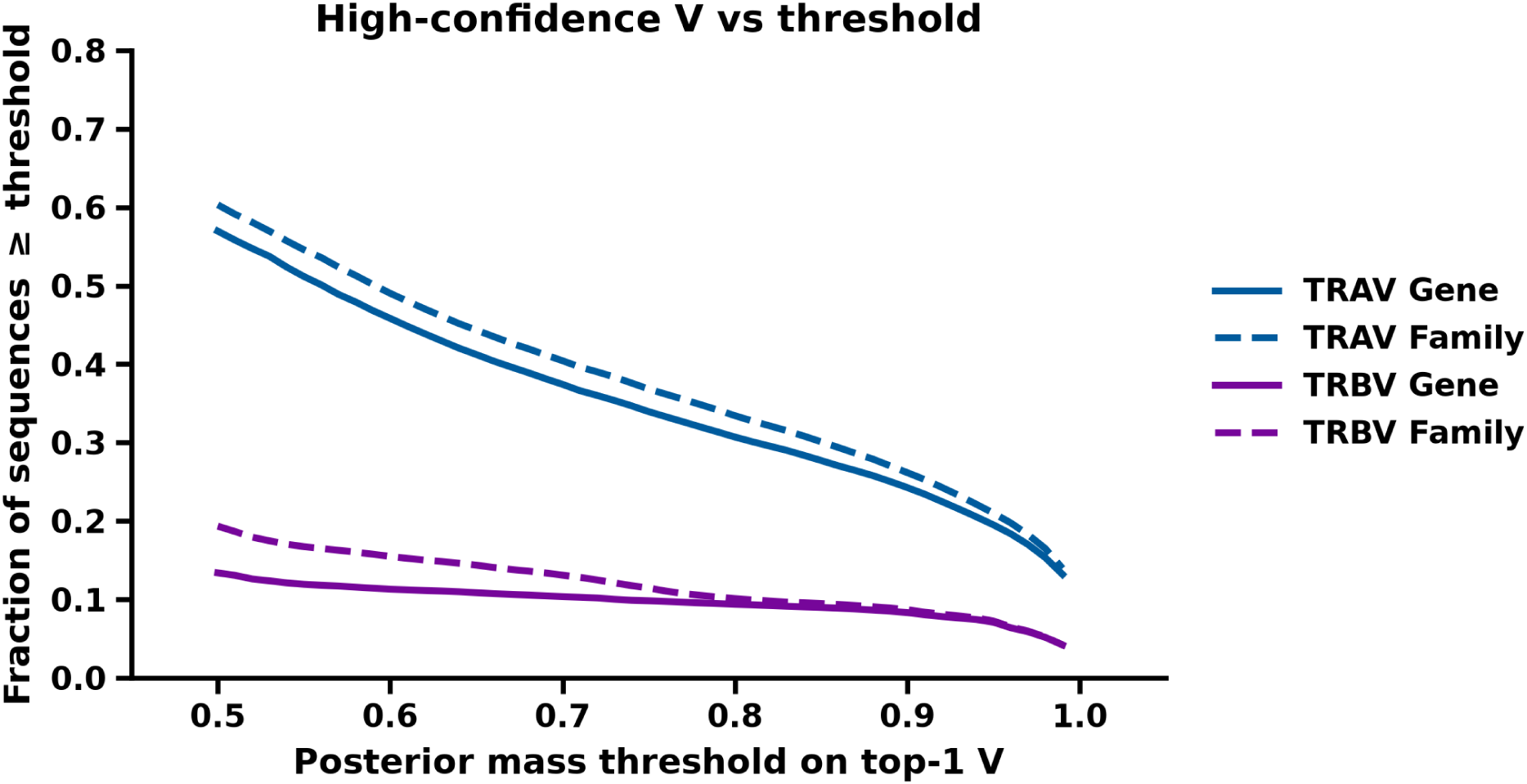
High-confidence single-gene V calls are attainable for only a minority. Fraction of sequences whose top-ranked V posterior mass meets a threshold (0.50..0.99), at gene and family resolution, TRA vs TRB. At the permissive 0.5 threshold only 13% of TRB sequences qualify at gene resolution (4% at 0.99); TRA reaches 57% (13% at 0.99). Coarsening to family barely lifts TRB (19% vs 13% at 0.5).

### 3.5 Selection sharpens the V posterior significantly but modestly

Post-selection sharpened the V posterior in both chains, but the effect, while statistically significant (*p <* 5.0 × 10*^−^*^4^ for both chains, the floor 1*/*(*n*_perm_ + 1) at 2,000 permutations, no permutation reaching either observed shift), was small in magnitude (Figure 6). After controlling for the change in marginal V usage, the mean V-gene conditional entropy shifted by −0.063 nats for *α*, −0.062 after correcting the small finite-sample estimator bias, and by −0.281 nats for *β*. The *α* raw shift was essentially zero (+0.0054), so the sharpening became visible only once usage was held fixed. Both shifts were significant against a label-permutation null, the *α* shift lying 36.9 and the *β* shift 206.5 standard deviations from it. *β* sharpened more than *α* at equal sample size, with the *α* shift falling outside the *β* matched-*n* interval [−0.2858, −0.2774] (Supplementary Figure S4). These shifts were small relative to the conditional entropies themselves, merely a fraction of a nat against pre-selection values of 1.3 and 2.7 nats, so selection refined the V posterior without rendering the individual V gene recoverable.

**Figure 6:**
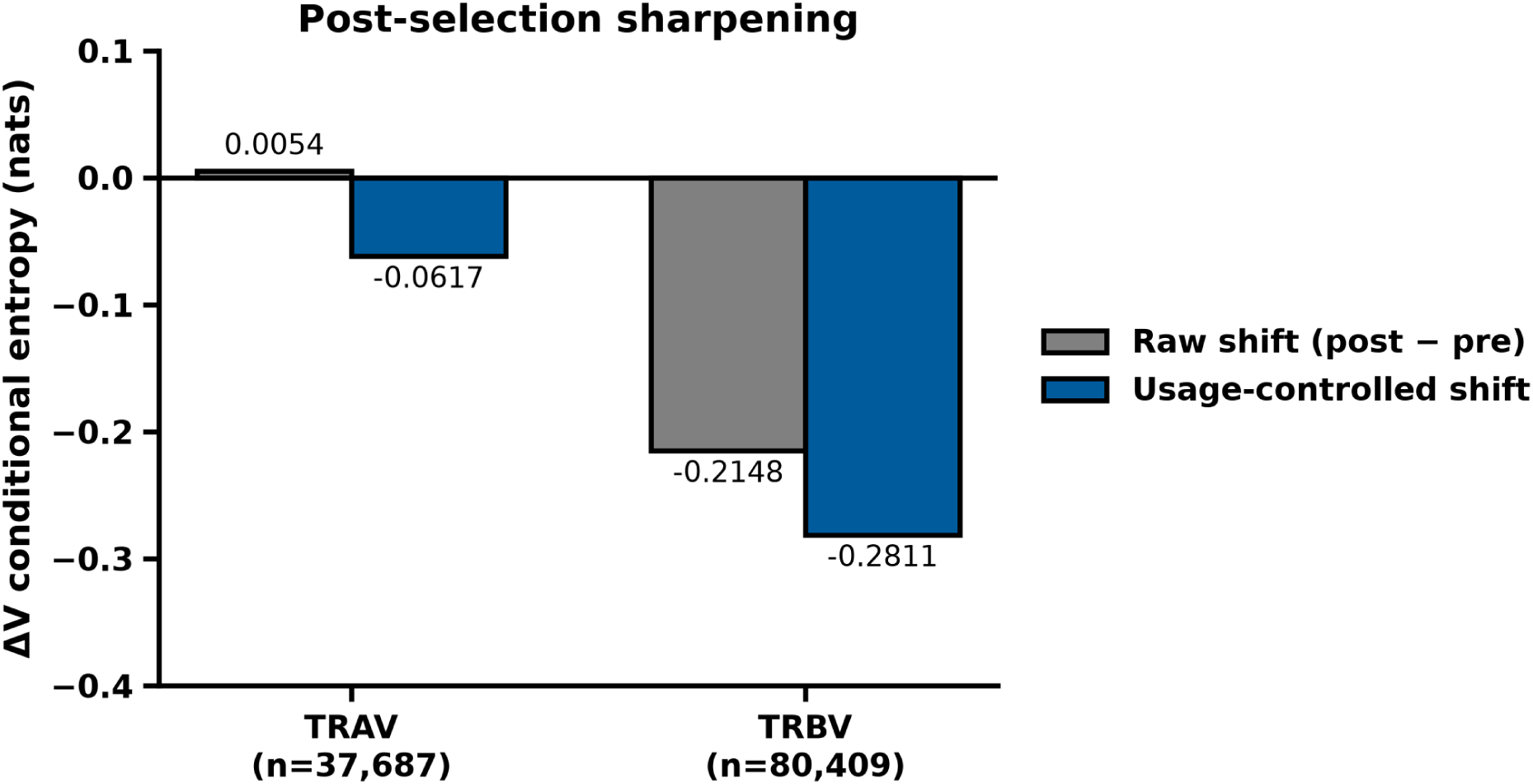
Selection sharpens the V posterior significantly but modestly. Raw and usage-controlled pre-to-post change in mean V-gene conditional entropy per chain; the TRA bar is additionally bias-corrected by the per-set permutation-null center. After controlling for the change in marginal V usage the shift is -0.063 nats for TRA (-0.062 bias-corrected) and -0.281 for TRB; the TRA raw shift is ∼0 (+0.0054), so sharpening emerges only once usage is held fixed.

### 3.6 Selection redistributes V identity, more substantially in ***β***

Beyond sharpening, selection changed which V gene was the most probable call, in the same direction in both chains. Suppression exceeded promotion, but with categorically different characteristics (Figure 7). In the *β* chain the redistribution was a substantial rewrite in which selection eliminated four genes as the argmax for any sequence, including TRBV13 (the top call for 15.97% of *β* sequences pre-selection) together with TRBV3-1, TRBV14, and TRBV24-1 (20,685 sequences in all, Supplementary Table S4). Six genes were promoted as the most probable V from zero pre-selection: dominantly TRBV2 (3,838 sequences), while amplification (promotion not necessarily from zero) was dominated by TRBV19 and TRBV4-1. The pre-selection argmax bin had no post-selection counterpart for 26.1% of *β* sequences against 7.2% in the reverse direction, and the dropped 7.2% concentrated almost entirely in TRBV2 and TRBV28 (Figure 8, Supplementary Table S5). In the *α* chain, the same operation was only a mild reweight: no gene appeared from zero and only a single TRAV18 sequence was eliminated as argmax; selection instead amplified already-present genes such as TRAV9-2 (from 1.38% to 9.13%), with the corresponding pre/post bin mismatches an order of magnitude smaller (0.40% and 0.15%). This contrast is consistent with the much larger *β* usage-controlled shift (−0.281 versus −0.063).

**Figure 7:**
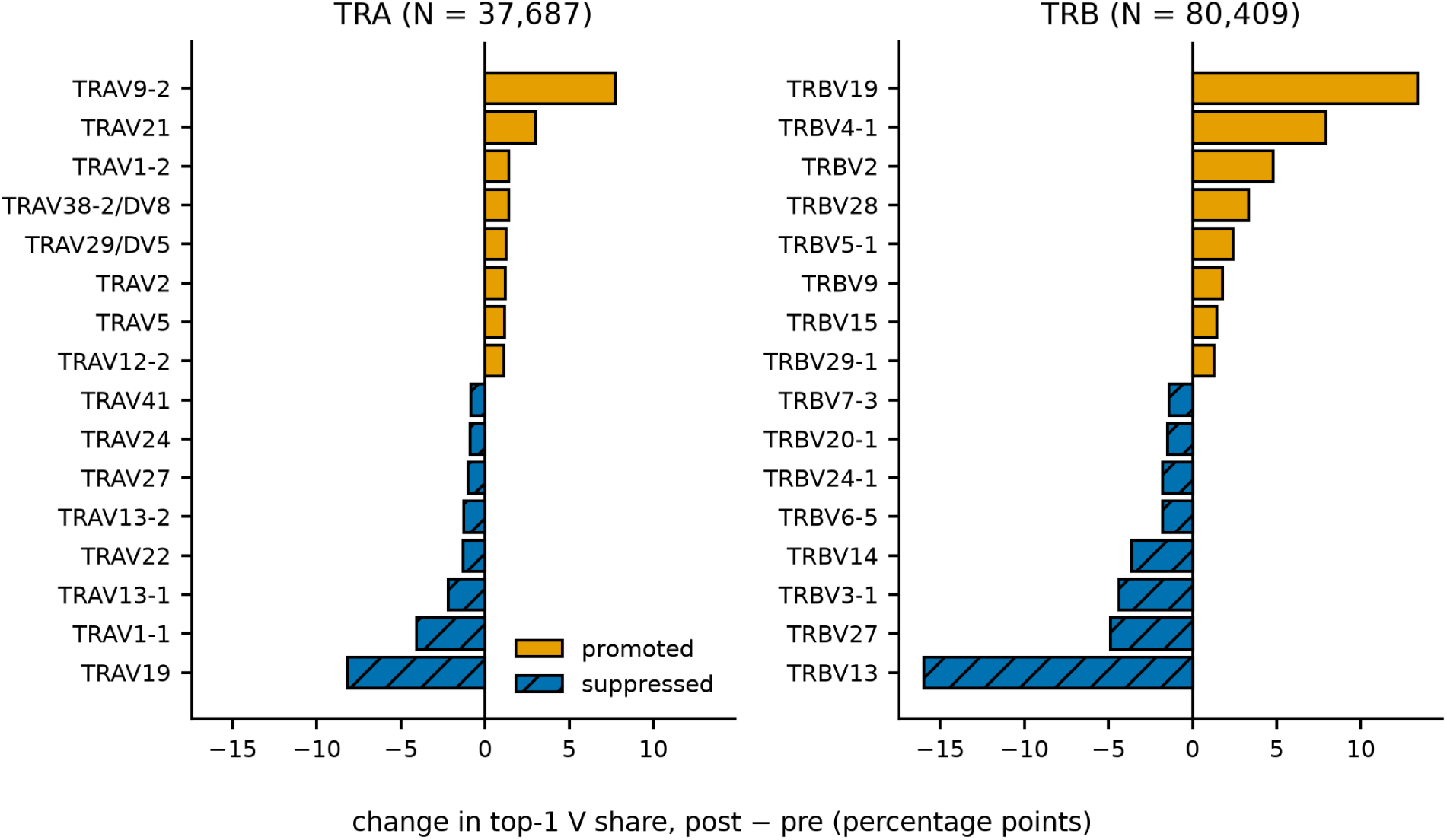
Selection redistributes the V argmax: a rewrite in beta, a reweight in alpha. Per-gene change in top-1 V share (post - pre) as diverging bars, **(A)** TRA and **(B)** TRB on a shared x-axis. In **(B)** selection eliminated four genes as the argmax (20,685 sequences, incl. TRBV13 at 15.97%) and created six from zero (dominantly TRBV2, 3,838); in **(A)** it only reweighted genes already present (e.g. TRAV9-2, 1.38% to 9.13%), erasing a single TRAV18 sequence.

**Figure 8:**
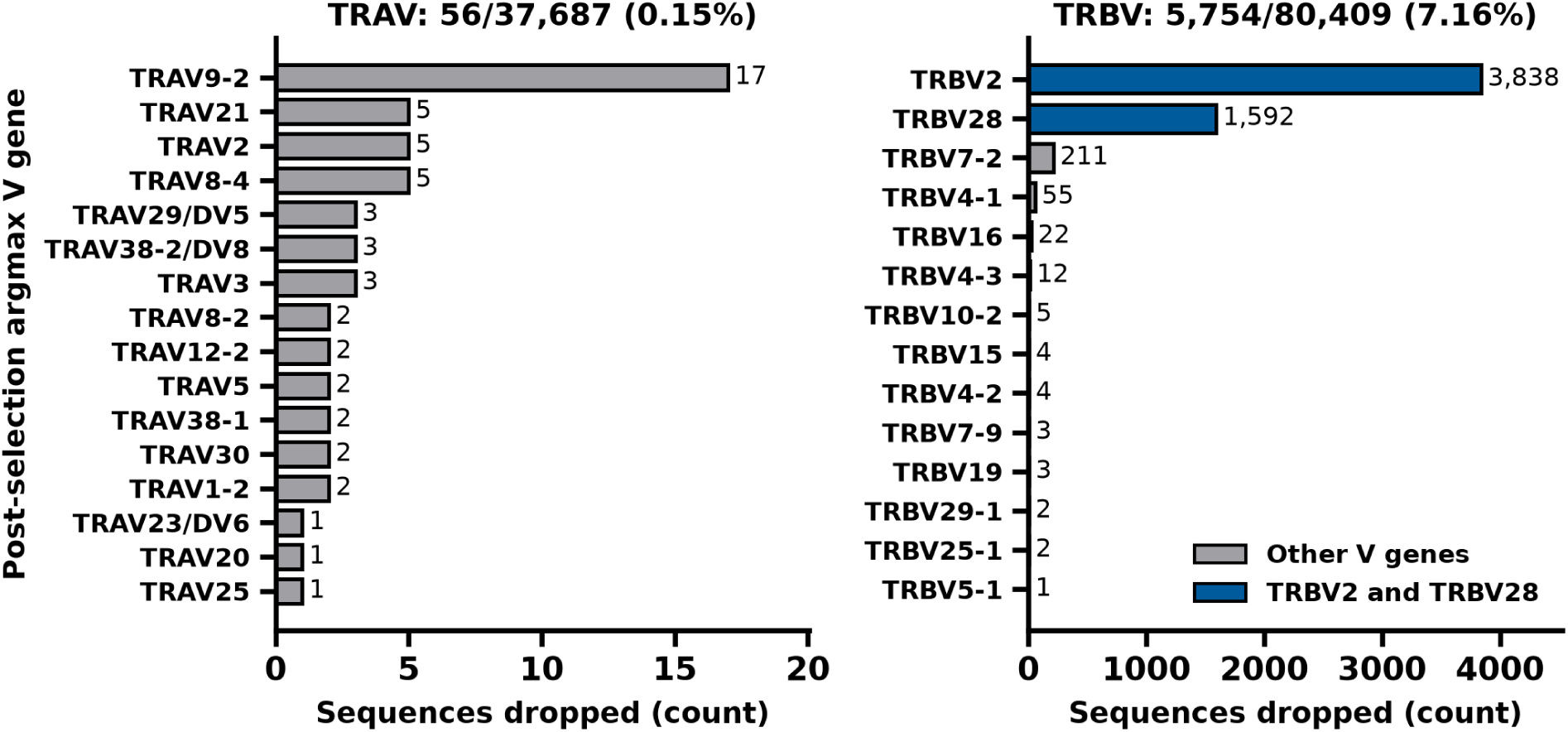
Where the usage control’s dropped weight concentrates, by chain. Per-gene count of sequences the usage control zeros (their CDR3-length x post-argmax-V cell is absent from the pre-selection marginal), **(A)** TRA and **(B)** TRB, independent x-axes. **(B)** drops 7.2% of the chain, 94% of it in TRBV2/TRBV28; **(A)** drops a negligible, diffuse 0.15% with no dominant gene.

### 3.7 The recoverability ceiling replicates in held-out tumor cohorts

The measurements were repeated on two held-out single-cell cohorts: a 231-patient atlas of anti-PD-1-treated non-small cell lung cancer [14, 15] and a 15-patient head and neck squamous cell carcinoma cohort treated with anti-PD-L1 and anti-CTLA-4 [16, 17], each reduced to held-out rearrangements and subsampled to 10,000 per cohort-chain (three cells, the head and neck cohort contributing *β* only) (Supplementary Table S6).

The *α* chain reproduced almost exactly. Mean V-gene conditional entropy was 1.2851 nats in the lung cohort against 1.2925 in the analysis set, the true V gene fell at rank one for 48% of sequences against 49% and within the top ten for 87% against 85%, and 57% of sequences met the 0.5 posterior-mass criterion in both. The *β* chain reproduced the qualitative result while sitting somewhat above the analysis set in absolute recoverability, with a V-gene entropy of 2.4365 nats in the lung cohort and 2.4565 in the head and neck cohort against 2.7055, the true V gene ranked first for 33% of sequences in both cohorts against 20%, and 23% against 13% of sequences meeting the 0.5 criterion.

That difference is attributable to V usage. Reweighting the analysis set so that its annotated-V marginal matched each cohort’s moved the *β* V-gene entropy from 2.7055 nats to 2.4181 (95% CI 2.4064 to 2.4299) against the lung cohort’s observed 2.4365, and to 2.4280 (2.4163 to 2.4400) against the head and neck cohort’s 2.4565. The reweighting therefore closes the whole of the 0.269 and 0.249 nat gaps and slightly overshoots them, leaving residuals of 0.018 and 0.029 nats in the opposite direction. Both cohorts are tumor-infiltrating repertoires dominated by clonally expanded T cells, and the narrower V usage that follows from that expansion accounts for the whole of their higher measured recoverability. The same reweighting applied to the *α* chain moved its entropy by 0.0045 nats against an observed gap of 0.0074.

The confusion structure itself also reappeared. Clustering each cohort’s own leakage matrix and comparing the resulting groups against the grouping derived from the analysis set gave an adjusted Rand index of 0.9594 for *α*, and 0.8671 and 0.8150 for *β* in the lung and head and neck cohorts, against 0.0519 and 0.214 for the agreement between the analysis-set grouping and the IMGT family partition. The genes that become mutually indistinguishable are therefore the same genes in both cohorts.

### 3.8 Bootstrap intervals and out-of-sample coverage

Resampling the 37,687 canonical *α* sequences with replacement over 1,000 replicates gave a 95% interval of [1.2834, 1.3013] nats on the *α* V-gene conditional entropy, [1.1764, 1.1931] at family resolution and [0.0106, 0.0123] on the J-gene entropy. The *β* V-gene entropy, bootstrapped at full sample size, fell in [2.6995, 2.7121]. The cross-chain gap is thus 1.4130 nats. The usage-controlled selection shift was −0.0632 nats with interval [−0.0740, −0.0563], or −0.0617 [−0.0726, −0.0549] once the finite-sample bias correction was applied to the point estimate and to both limits; the interval excludes zero, agreeing with the permutation test. Re-deriving the V confusion grouping inside each replicate and scoring it against the full-data grouping gave a mean adjusted Rand index of 0.944 [0.844, 1.000], so the groups in Figure 3 are a property of the *α* set rather than of one particular sample of it.

The candidate-set utility was then checked on data it had not seen. The coverage curve was fitted on half the canonical set and tested on the other half, with sequences assigned to halves by a hash of the CDR3. Across both chains and every level from 0.5 to 0.95, requested and achieved coverage differed by at most 0.4 percentage points. The largest gap was 0.0039, and asking for 0.95 returned 0.9486 in each chain.

## 4 Discussion

The recoverability of V and J genes from a CDR3 sequence alone is limited and asymmetric between the two genes. The J gene was nearly perfectly recovered by the CDR3 in both chains, so recovering J from a CDR3 is for practical purposes a solved problem, although the strong germline presence of J is already well understood. The V gene, by contrast, was only partially recoverable, and substantially more so for the *α* chain than for the *β* chain across the full range of CDR3 lengths. The residual V uncertainty was not random scatter around a single correct gene but structured confusion among groups of genes, and those groups did not overlap with the IMGT gene families. The natural unit of V recovery is therefore a group of mutually confusable genes that crosses IMGT family boundaries, recoverable where the individual gene is not, but not aligned with the nomenclatural family.

This group structure is an emergent property of the repertoire rather than a feature of any one gene or any one receptor. It has no representation in a single rearrangement: it is defined only by how posterior mass is distributed across the entire V locus, and it appeared here only because 118,096 rearrangements were read jointly. It is also not deducible from the germline sequences, which is why the data-driven grouping departs so far from the IMGT family partition (adjusted Rand index 0.0519 for *α* and 0.214 for *β*); a nomenclature built from germline similarity does not predict which genes become indistinguishable once the junction is written. The structure instead arises from processes acting at different levels of the same system: exonucleolytic trimming and non-templated insertion erase germline V identity at the junction, translation to amino acids discards the synonymous codon information that would otherwise separate similar genes, and the selection filter then reweights which genes carry the residual posterior mass.

This reframing carries an implication for tools that assign V identity from a CDR3 alone. Because the recoverable object is a group instead of a gene, such tools should return a calibrated candidate set, meaning a confusion group with an associated confidence. The confidence curves reinforce this for the *β* chain, where a high-confidence single-gene call was available for only a small minority of sequences even at a permissive threshold. The *α* chain is substantially more favorable but still limited, so reporting a group with calibrated confidence is a safe method for both chains.

Comparing the pre-selection recombination posterior with the post-selection posterior, the molecular machinery that generates the repertoire against the organism-level filter that shapes what survives in it revealed a second, smaller effect. Selection modestly sharpened the V posterior in both chains beyond any change in marginal usage. The sharpening was statistically significant in both chains and stronger in *β* than in *α*, yet it was relatively small compared to the conditional entropies themselves and far from sufficient to consider the V gene recoverable. Selection also redistributed which V gene was the most probable call, with different characteristics between the chains. In the *β* chain, selection rewrote the argmax landscape, eliminating several genes as the top call for any sequence and promoting others from zero, whereas in *α* selection merely reweighted genes already present, amplifying some without eliminating anything material.

This redistribution should be weighed against experimental analyses of pre- and post-selection repertoires, which compare modeled or thymocyte baselines against naive cells and report only a weak effect of selection on V and J gene usage [22]. Our redistribution is a model-attributed reshaping of the argmax call rather than a measured change in gene-usage frequency. It describes which V genes the pretrained SONNIA model favors and disfavors, and whether its cause is biological or a property of the model’s training data is not established here.

Several limitations bound these conclusions. The recoverability ceiling is defined relative to the OLGA and SONNIA generative models. This study quantifies the information a CDR3 carries about V and J under those models, so the ceiling is model-based rather than absolute and a different generative model could shift it. Clonal similarity reduces the effective sample size, so the permutation p-values are lower bounds. The selection findings carry a further point of discussion, being that both the merged dataset and the cohort on which the SONNIA models were trained are enriched for viral-responsive, antigen-experienced TCRs, so part of the apparent selection signal may reflect shared viral-repertoire composition between the data and the selection model rather than selection alone. This confound would inflate the measured effect rather than create the structured, gene-level redistribution we observe, and the sign of the selection shift was stable across the datasets despite their differing composition. Nonetheless, separating selection from shared repertoire composition would require comparison against an unselected in-vivo baseline, such as non-productive rearrangements, which is beyond the scope of this analysis. Finally, all three databases are epitope-associated and antigen-sorted, so these conclusions may not generalize to a bulk, unsorted repertoire. The ceiling itself, however, is not specific to those databases: both the recoverability measurements and the confusion grouping reproduced in two tumor cohorts that contributed nothing to the analysis set, with the *α* V-gene entropy within 0.008 nats and the grouping agreeing at an adjusted Rand index of 0.96. Those cohorts are themselves clonally expanded tumor-infiltrating repertoires rather than bulk unsorted ones, so they place the result beyond the source databases without settling the bulk case. In sum, this work contributes a quantified identifiability ceiling for inferring V and J genes from TCR CDR3 sequences, together with a statement of what is recoverable: J almost completely, and V only as a data-driven confusion group whose resolution is higher for *α* than for *β* and does not follow IMGT family boundaries. For anyone inferring V or J from a CDR3, in repertoire annotation, single-cell reconstruction, or computation of missing fields in TCR repertoire data, these results delineate what can and cannot be recovered and argue for reporting calibrated groups rather than single genes.

## Abbreviations

ARI: adjusted Rand index;
CDR3: third complementarity-determining region;
CI: confidence interval;
GEO: Gene Expression Omnibus;
HNSCC: head and neck squamous cell carcinoma;
IMGT: the international ImMunoGeneTics information system;
NSCLC: non-small cell lung cancer;
*P*_gen_: generation probability;
*P*_post_: post-selection probability;
TCR: T-cell receptor;
TRA/TRB: TCR *α*/*β* chain;
TRAV/TRBV: TRA/TRB variable gene;
TRAJ/TRBJ: TRA/TRB joining gene;
V(D)J: variable–(diversity)–joining.

## Ethics Statement

Ethical approval was not required for this study in accordance with the local legislation and institutional requirements. The study analyzes only data that are publicly available, previously published and de-identified: T-cell receptor sequences deposited in VDJdb, McPAS-TCR and NeoTCR, and the two published single-cell cohorts obtained from the Gene Expression Omnibus. No new human subjects research was undertaken, no new samples were collected, and no identifiable participant information was accessed.

## Author Contributions

SJH: Conceptualization, Data curation, Formal analysis, Investigation, Methodology, Software, Validation, Visualization, Writing – original draft, Writing – review & editing. ASB: Supervision, Resources, Writing – review & editing.

## Generative AI Statement

The author(s) declare that generative AI was used in the creation of this manuscript. The author(s) declare that Claude Opus 5 was used for editing and formatting.

## Supplementary Material

Supplementary Figures S1–S4 and Supplementary Tables S1–S6 are provided as a separate Supplementary Material file. The per-gene tables are additionally supplied as CSV files under results/supp tables/ in the deposited data record, https://doi.org/10.5281/zenodo.22024426.

## Data Availability Statement

The datasets analyzed for this study are public. The repertoire sources are VDJdb (https://vdjdb.com/), McPAS-TCR (https://friedmanlab.weizmann.ac.il/McPAS-TCR/), and NeoTCR (http://neotcrdb.bioxai.cn/). The held-out validation cohorts are in the Gene Expression Omnibus under accessions GSE243013 and GSE286827. The analysis pipeline, the calibrated candidate-set utility and the scripts that regenerate every number and figure in this manuscript are at https://github.com/samhuann/SuperVDJMachine, archived at https://doi.org/10.5281/ zenodo.22025548 [23]. The per-sequence posterior table and the analysis artifacts from which every figure and reported value is derived are deposited at https://doi.org/10.5281/zenodo.22024426 [24]. All figures are plotted directly from the deposited posterior table by those scripts; no image processing was applied to any figure.

## References

[1] Krangel MS. Mechanics of T cell receptor gene rearrangement. Curr Opin Immunol (2009) 21:133–9. doi: 10.1016/j.coi.2009.03.009

[2] Lefranc MP. From IMGT-ONTOLOGY CLASSIFICATION Axiom to IMGT Standardized Gene and Allele Nomenclature: For Immunoglobulins (IG) and T Cell Receptors (TR). Cold Spring Harb Protoc (2011) 2011:pdb.ip84. doi: 10.1101/pdb.ip84

[3] Lefranc MP, Pommíe C, Ruiz M, Giudicelli V, Foulquier E, Truong L, et al. IMGT unique numbering for immunoglobulin and T cell receptor variable domains and Ig superfamily V-like domains. Dev Comp Immunol (2003) 27:55–77. doi: 10.1016/s0145-305x(02)00039-3

[4] Shugay M, Bagaev DV, Zvyagin IV, Vroomans RM, Crawford JC, Dolton G, et al. VDJdb: a curated database of T-cell receptor sequences with known antigen specificity. Nucleic Acids Res (2018) 46:D419–27. doi: 10.1093/nar/gkx760

[5] Gerritsen B, Pandit A, Andeweg AC, de Boer RJ. RTCR: a pipeline for complete and accurate recovery of T cell repertoires from high throughput sequencing data. Bioinformatics (2016) 32:3098–106. doi: 10.1093/bioinformatics/btw339

[6] Springer I, Tickotsky N, Louzoun Y. Contribution of T Cell Receptor Alpha and Beta CDR3, MHC Typing, V and J Genes to Peptide Binding Prediction. Front Immunol (2021) 12:664514. doi: 10.3389/fimmu.2021.664514

[7] Sidhom JW, Larman HB, Pardoll DM, Baras AS. DeepTCR is a deep learning framework for revealing sequence concepts within T-cell repertoires. Nat Commun (2021) 12:1605. doi: 10.1038/s41467-021-21879-w

[8] Dash P, Fiore-Gartland AJ, Hertz T, Wang GC, Sharma S, Souquette A, et al. Quantifiable predictive features define epitope-specific T cell receptor repertoires. Nature (2017) 547:89–93. doi: 10.1038/nature22383

[9] Leary AY, Scott D, Gupta NT, Waite JC, Skokos D, Atwal GS, et al. Designing meaningful continuous representations of T cell receptor sequences with deep generative models. Nat Commun (2024) 15:4271. doi: 10.1038/s41467-024-48198-0

[10] Moreno DL, Croce G, Gfeller D. Statistical modelling of CDR3 sequences provides robust quality control for TCR repertoire datasets. bioRxiv [Preprint] (2025). Available at: 10.64898/2025.12.11.693618 (Accessed August 20, 2026)

[11] Sethna Z, Elhanati Y, Callan CG, Walczak AM, Mora T. OLGA: fast computation of generation probabilities of B- and T-cell receptor amino acid sequences and motifs. Bioinformatics (2019) 35:2974–81. doi: 10.1093/bioinformatics/btz035

[12] Marcou Q, Mora T, Walczak AM. High-throughput immune repertoire analysis with IGoR. Nat Commun (2018) 9:561. doi: 10.1038/s41467-018-02832-w

[13] Isacchini G, Walczak AM, Mora T, Nourmohammad A. Deep generative selection models of T and B cell receptor repertoires with soNNia. Proc Natl Acad Sci U S A (2021) 118:e2023141118. doi: 10.1073/pnas.2023141118

[14] Liu Z, Yang Z, Wu J, Zhang W, Sun Y, Zhang C, et al. A single-cell atlas reveals immune heterogeneity in anti-PD-1-treated non-small cell lung cancer. Cell (2025) 188:3081–3096.e19. doi: 10.1016/j.cell.2025.03.018

[15] Liu Z, Yang Z, Wu J, Zhang W, Sun Y, Zhang C, et al. Single-cell RNA and TCR sequencing of anti-PD-1-treated non-small cell lung cancer [Dataset]. Gene Expression Omnibus (2025). Accession: GSE243013. Available at: https://www.ncbi.nlm.nih.gov/geo/query/acc.cgi?acc=GSE243013

[16] Cha J, Kim CG, Sim NS, Kim G, Son W, Kim D, et al. 4-1BB^+^ Tregs and inhibitory progenitor exhausted T cells confer resistance to anti-PD-L1 and anti-CTLA-4 combination therapy. Cell Rep Med (2025) 6:102408. doi: 10.1016/j.xcrm.2025.102408

[17] Cha J, Kim CG, Sim NS, Kim G, Son W, Kim D, et al. Single-cell profiling of head and neck squamous cell carcinoma under anti-PD-L1 and anti-CTLA-4 therapy [Dataset]. Gene Expression Omnibus (2025). Accession: GSE286827. Available at: https://www.ncbi.nlm.nih.gov/geo/query/acc.cgi?acc=GSE286827

[18] Zhou W, Xiang W, Yu J, Ruan Z, Pan Y, Wang K, et al. NeoTCR: An Immunoinformatic Database of Experimentally-supported Functional Neoantigen-specific TCR Sequences. Genomics Proteomics Bioinformatics (2025) 23:qzae010. doi: 10.1093/gpbjnl/qzae010

[19] Tickotsky N, Sagiv T, Prilusky J, Shifrut E, Friedman N. McPAS-TCR: a manually curated catalogue of pathology-associated T cell receptor sequences. Bioinformatics (2017) 33:2924–9. doi: 10.1093/bioinformatics/btx286

[20] Carlson CS, Emerson RO, Sherwood AM, Desmarais C, Chung MW, Parsons JM, et al. Using synthetic templates to design an unbiased multiplex PCR assay. Nat Commun (2013) 4:2680. doi: 10.1038/ncomms3680

[21] Giudicelli V, Chaume D, Lefranc MP. IMGT/GENE-DB: a comprehensive database for human and mouse immunoglobulin and T cell receptor genes. Nucleic Acids Res (2005) 33:D256–61. doi: 10.1093/nar/gki010

[22] Luppov DV, Vlasova EK, Chudakov DM, Shugay M. Comprehensive analysis of *αβ* T-cell receptor repertoires reveals signatures of thymic selection. Front Immunol (2025) 16:1605170. doi: 10.3389/fimmu.2025.1605170

[23] Huang SJ, Baras AS. SuperVDJMachine: posteriors over TCR V and J genes from CDR3 sequences [Software]. Zenodo (2026). Version v1. Available at: 10.5281/zenodo.22025548

[24] Huang SJ, Baras AS. Per-sequence V and J posterior table and analysis artifacts for: Quantifying the Recoverability of V and J Genes from TCR CDR3 Sequences Using Generative Repertoire Models [Dataset]. Zenodo (2026). Version v1. Available at: 10.5281/zenodo.22024426

