## Supplementary Material for "Quantifying the Recoverability of V and J Genes from TCR CDR3 Sequences Using Generative Repertoire Models"

### 1 Supplementary Figures and Tables

This file accompanies the associated manuscript and contains Supplementary Figures S1–S4 and Supplementary Tables S1–S6. The per-gene tables behind them are supplied as CSV files in the deposited data record, `https://doi.org/10.5281/zenodo.22024426`.

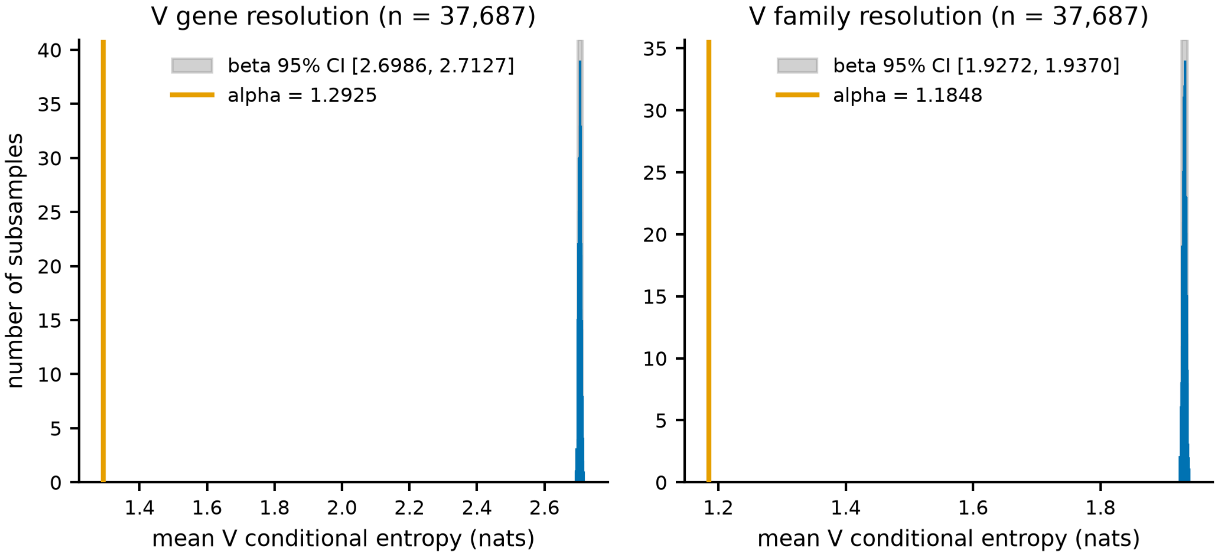


**Supplementary Figure S1.** Alpha V-entropy lies below beta at matched sample size. TRB subsampled without replacement to the TRA sample size (n=37,687) over 500 replicates; histogram = beta matched-n mean V conditional entropy, orange line = alpha value, grey band = beta 95% CI. **(A)** Gene resolution: alpha 1.2925 vs beta CI [2.6986, 2.7127]. **(B)** Family resolution: alpha 1.1848 vs [1.9272, 1.9370]. Alpha lies far below every beta draw in both panels.

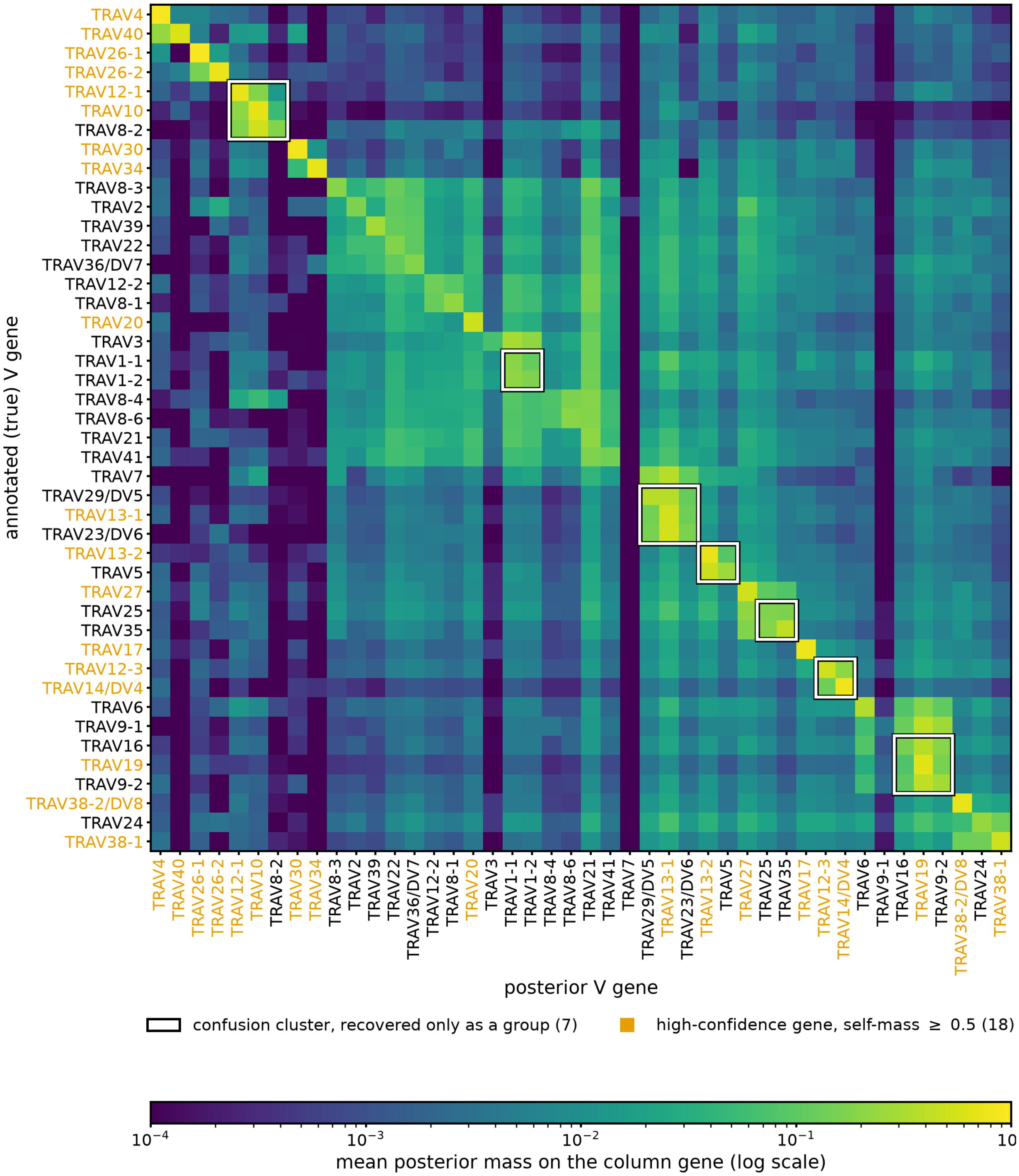


**Supplementary Figure S2.** TRAV confusion matrix annotated with recovery structure. White boxes mark the 7 data-driven confusion clusters of $\geq 2$ genes; genes inside a box are recoverable only as a group. Orange gene labels mark the 18 high-confidence recovery genes whose mean posterior self-mass (matrix diagonal) is $\geq 0.5$, i.e. the correct gene receives the majority of the posterior on average and can be called individually. Row = true V gene, column = posterior V gene; cell = mean posterior mass (log scale). V genes with <20 sequences omitted.

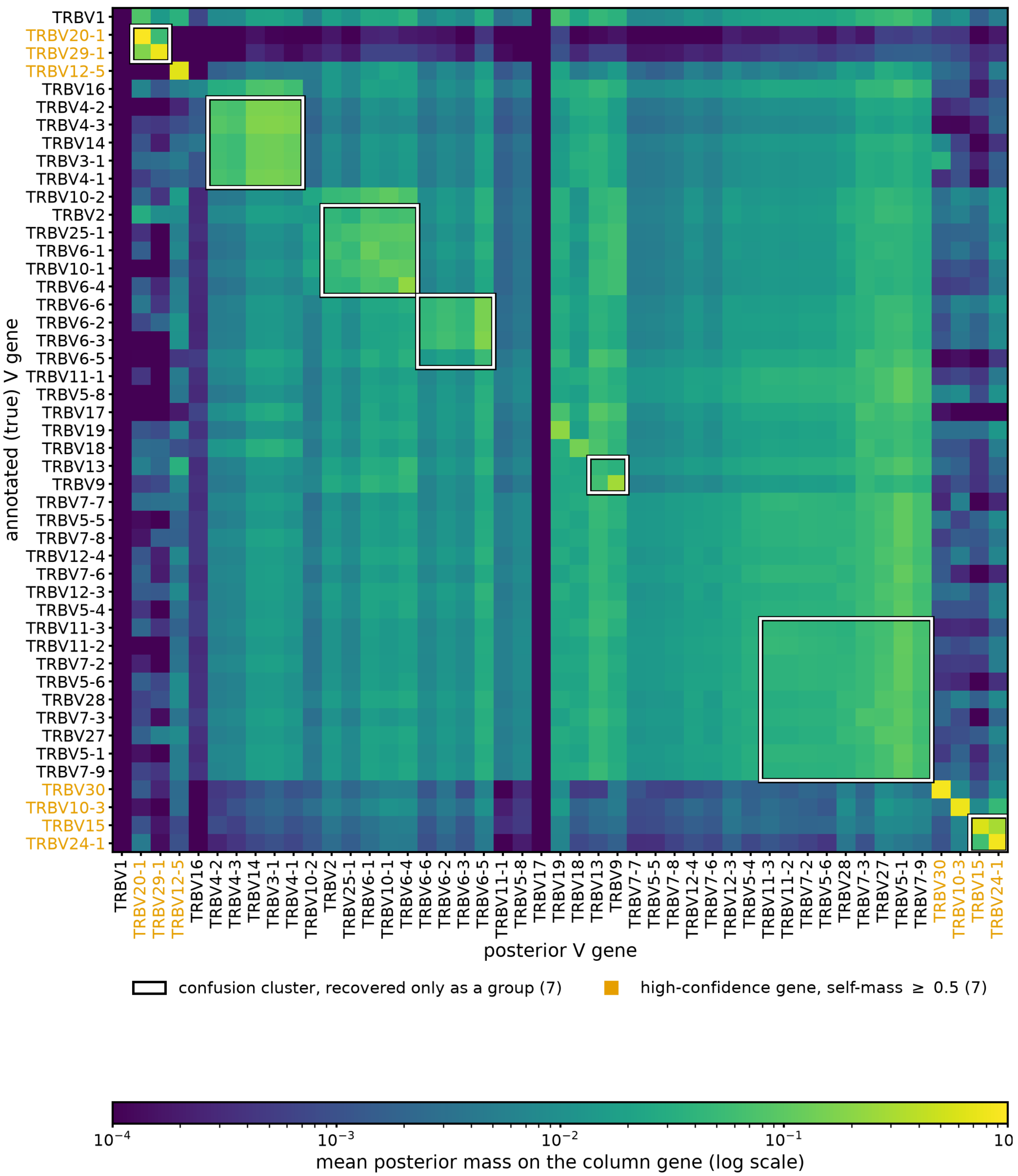


**Supplementary Figure S3.** TRBV confusion matrix annotated with recovery structure. White boxes mark the 7 data-driven confusion clusters of $\geq 2$ genes; genes inside a box are recoverable only as a group. Orange gene labels mark the 7 high-confidence recovery genes whose mean posterior self-mass (matrix diagonal) is $\geq 0.5$, i.e. the correct gene receives the majority of the posterior on average and can be called individually. Row = true V gene, column = posterior V gene; cell = mean posterior mass (log scale). V genes with <20 sequences omitted.

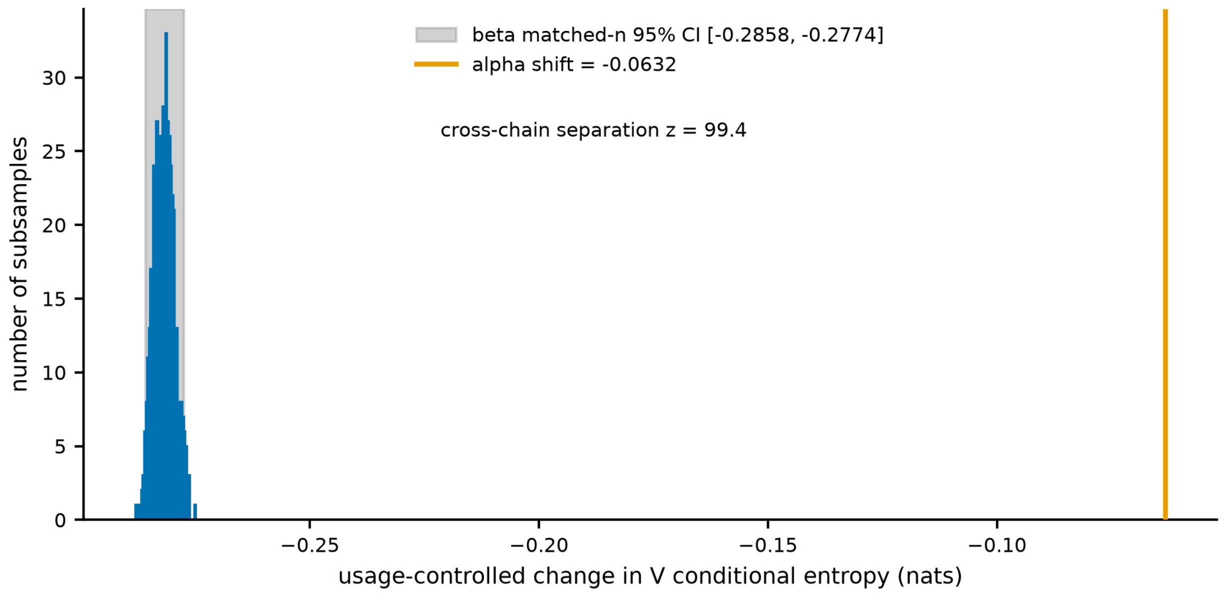


**Supplementary Figure S4.** Beta sharpens more than alpha at matched sample size. Usage-controlled V selection shift with TRB subsampled to the TRA n=37,687 over 500 replicates (histogram = beta matched-n shift, mean -0.2819, 95% CI [-0.2858, -0.2774]; orange line = alpha shift -0.063, the uncorrected usage-controlled value; the bias-corrected -0.0617 in Figure 6 of the main text subtracts the permutation-null center). Alpha sits outside the beta interval, beyond every beta draw (cross-chain separation z = 99.4 sigma).

**Supplementary Table S1.** Schema of the full posterior table, deposited at `https://doi.org/10.5281/zenodo.22024426` [reference 25].

| **column** | **description** |
|---|---|
| `seq_id` | identifier of the canonical rearrangement |
| `chain` | TRA / TRB |
| `cdr3` | CDR3 amino-acid sequence |
| `axis` | V / J — which gene the posterior is over |
| `model` | pre (OLGA) / post (OLGA x SONNIA selection) |
| `mode` | grid (full V x J) / fixed (condition on annotated J; present in the deposited table but not used for any analysis reported here) / NaN for pre |
| `resolution` | gene / family |
| `status` | ok / impossible (OLGA $P_{\text{gen}}$=0 → empty posterior) |
| `entropy_nats` | per-sequence posterior conditional entropy (nats); NaN if empty |
| `top1_mass` | maximum posterior mass; NaN if empty |
| `top1_label` | argmax gene/family label |
| `posterior_json` | full posterior distribution as JSON; '{}' if empty |

**Supplementary Table S2.** Three-way $\alpha$ source robustness. Rows: $\alpha$ analyzed on McPAS alone, VDJdb alone, and the pooled set. Columns: n (canonical sequences, i.e. status "ok" after the zero-$P_{\text{gen}}$ sequences are removed; the corresponding pre-canonical intra-source counts are 6,118, 33,081 and 38,971); v_gene_entropy (mean V-gene conditional entropy, nats); ctrl_selection_shift (usage-controlled pre-to-post entropy shift, nats); grouping_ari (adjusted Rand index of the data-driven V confusion grouping vs IMGT families). The intra-source canonical counts do not sum to the pooled row because 486 canonical rearrangements occur in more than one source, one of them in all three; removing the 487 duplicate memberships gives $5{,}829 + 32{,}085 + 260 - 487 = 37{,}687$, where 260 is the canonical NeoTCR-$\alpha$ count. NeoTCR-$\alpha$ is not shown as a standalone row: it contributes only 267 chains (260 canonical), spread over 41 V genes; those sequences are included in the pooled row.

| **source** | **n** | **v_gene_entropy** | **ctrl_selection_shift** | **grouping_ari** |
|---|---|---|---|---|
| McPAS-alpha | 5,829 | 1.334 | −0.093 | 0.062 |
| VDJdb-alpha | 32,085 | 1.291 | −0.061 | 0.060 |
| Pooled-alpha | 37,687 | 1.293 | −0.063 | 0.052 |

**Supplementary Table S3.** High-confidence V-gene recovery. Genes whose mean posterior self-mass (the confusion-matrix diagonal: the average posterior probability placed on the correct gene) is $\geq 0.50$, so the annotated gene receives the majority of the posterior on average and can be called individually from the CDR3. Columns: chain (TRA/TRB); gene (IMGT V gene); self_mass (mean posterior self-mass, canonical set, gene resolution). 18 alpha and 7 beta genes qualify; sorted by self_mass within chain. These are the orange-labelled genes in Supplementary Figures S2 and S3.

**V$\alpha$ (n=18)**

| **gene** | **self-mass** |
|---|---|
| TRAV26-1 | 0.9131 |
| TRAV4 | 0.9111 |
| TRAV30 | 0.8288 |
| TRAV14/DV4 | 0.7956 |
| TRAV17 | 0.7926 |
| TRAV26-2 | 0.7585 |
| TRAV13-2 | 0.7258 |
| TRAV38-2/DV8 | 0.7187 |
| TRAV34 | 0.7136 |
| TRAV12-1 | 0.7119 |
| TRAV10 | 0.6916 |
| TRAV19 | 0.6624 |
| TRAV40 | 0.5956 |
| TRAV13-1 | 0.5418 |
| TRAV38-1 | 0.5201 |
| TRAV12-3 | 0.5146 |
| TRAV27 | 0.5071 |
| TRAV20 | 0.5011 |

**V$\beta$ (n=7)**

| **gene** | **self-mass** |
|---|---|
| TRBV20-1 | 0.9368 |
| TRBV30 | 0.8826 |
| TRBV24-1 | 0.8611 |
| TRBV29-1 | 0.8008 |
| TRBV10-3 | 0.7823 |
| TRBV12-5 | 0.6496 |
| TRBV15 | 0.5873 |

**Supplementary Table S4.** Per-gene selection redistribution of the argmax V call (canonical set, gene resolution). Columns: chain; gene; category, coded E, P or A (E = eliminated as argmax: pre-selection top-1 count > 0 but post-selection top-1 count = 0; P = promoted from zero: pre top-1 = 0 but post top-1 > 0; A = amplified: present in both, largest positive change in top-1 share); pre_top1/post_top1 = number of sequences for which the gene is the highest-posterior V pre-/post-selection; pre_top1_share/post_top1_share = those counts / N; the per-gene marginal usage columns (mean posterior mass pre- and post-selection) are in the accompanying CSV rather than here; delta_top1_share = post_top1_share - pre_top1_share; shares use the canonical N for the chain (37,687 $\alpha$, 80,409 $\beta$). Companion table `results/supp_tables/tableS4_redistribution_accounting.csv` in the deposited data record [reference 25]: promotion = post-argmax (CDR3-length x gene) cells absent from the pre marginal (these sequences are dropped by the usage control); suppression = pre-argmax cells absent post (suppression_kept = suppressed sequences retained because their post cell exists in pre); bins = distinct (length x gene) cells; dropped_total equals promotion_seqs.

| chain | gene | category | pre_top1 | post_top1 | pre_top1_share | post_top1_share | delta_top1_share |
|---|---|---|---|---|---|---|---|
| TRB | TRBV2 | P | 0 | 3,838 | 0 | 0.0477 | 0.0477 |
| TRB | TRBV7-2 | P | 0 | 211 | 0 | 0.0026 | 0.0026 |
| TRB | TRBV16 | P | 0 | 22 | 0 | 0.0003 | 0.0003 |
| TRB | TRBV4-3 | P | 0 | 12 | 0 | 0.0001 | 0.0001 |
| TRB | TRBV10-2 | P | 0 | 5 | 0 | 0.0001 | 0.0001 |
| TRB | TRBV4-2 | P | 0 | 4 | 0 | 0 | 0 |
| TRB | TRBV19 | A | 3,294 | 14,063 | 0.041 | 0.1749 | 0.1339 |
| TRB | TRBV4-1 | A | 683 | 7,070 | 0.0085 | 0.0879 | 0.0794 |
| TRB | TRBV28 | A | 22 | 2,684 | 0.0003 | 0.0334 | 0.0331 |
| TRB | TRBV5-1 | A | 19,214 | 21,147 | 0.239 | 0.263 | 0.024 |
| TRB | TRBV9 | A | 7,483 | 8,900 | 0.0931 | 0.1107 | 0.0176 |
| TRB | TRBV13 | E | 12,845 | 0 | 0.1597 | 0 | −0.1597 |
| TRB | TRBV3-1 | E | 3,508 | 0 | 0.0436 | 0 | −0.0436 |
| TRB | TRBV14 | E | 2,898 | 0 | 0.036 | 0 | −0.036 |
| TRB | TRBV24-1 | E | 1,434 | 0 | 0.0178 | 0 | −0.0178 |
| TRA | TRAV9-2 | A | 520 | 3,441 | 0.0138 | 0.0913 | 0.0775 |
| TRA | TRAV21 | A | 4,044 | 5,182 | 0.1073 | 0.1375 | 0.0302 |
| TRA | TRAV1-2 | A | 801 | 1,333 | 0.0213 | 0.0354 | 0.0141 |
| TRA | TRAV38-2/DV8 | A | 1,127 | 1,657 | 0.0299 | 0.044 | 0.0141 |
| TRA | TRAV29/DV5 | A | 780 | 1,257 | 0.0207 | 0.0334 | 0.0127 |
| TRA | TRAV18 | E | 1 | 0 | 0 | 0 | −0 |

| chain | n | promotion_bins | promotion_seqs | promotion_pct | suppression_bins | suppression_seqs | suppression_pct | suppression_kept |
|---|---|---|---|---|---|---|---|---|
| TRB | 80,409 | 64 | 5,754 | 7.156 | 81 | 20,997 | 26.113 | 20,564 |
| TRA | 37,687 | 26 | 56 | 0.149 | 31 | 149 | 0.395 | 122 |

**Supplementary Table S5.** Usage-control dropped-weight concentration ($\beta$). Columns: gene (post-selection argmax V); dropped (sequences the usage control zeros because their (CDR3-length x argmax-V) cell is absent from the pre marginal); pct_of_dropped (share of all dropped weight); gene_drop_rate (dropped / total post-argmax sequences for that gene); post_top1_frac_of_chain (gene's share of all $\beta$ post-argmax calls). Summary: 5,754 / 80,409 (7.2%) dropped, 94.4% in TRBV2+TRBV28 (66.7% + 27.7%); drop rate 100% TRBV2, 59.3% TRBV28. Smoothing robustness: retaining the dropped sequences via an additively smoothed pre marginal moves the controlled shift only from -0.2811 to -0.2830 nats.

| **gene** | **dropped** | **pct_of_dropped** | **gene_drop_rate** | **post_top1_frac_of_chain** |
|---|---:|---:|---:|---:|
| TRBV2 | 3,838 | 66.7 | 100 | 4.773 |
| TRBV28 | 1,592 | 27.67 | 59.31 | 3.338 |
| TRBV7-2 | 211 | 3.67 | 100 | 0.262 |
| TRBV4-1 | 55 | 0.96 | 0.78 | 8.793 |
| TRBV16 | 22 | 0.38 | 100 | 0.027 |
| TRBV4-3 | 12 | 0.21 | 100 | 0.015 |
| TRBV10-2 | 5 | 0.09 | 100 | 0.006 |
| TRBV15 | 4 | 0.07 | 0.24 | 2.079 |
| TRBV4-2 | 4 | 0.07 | 100 | 0.005 |
| TRBV7-9 | 3 | 0.05 | 3.19 | 0.117 |
| TRBV19 | 3 | 0.05 | 0.02 | 17.489 |
| TRBV29-1 | 2 | 0.03 | 0.08 | 3.184 |
| TRBV25-1 | 2 | 0.03 | 0.71 | 0.349 |
| TRBV5-1 | 1 | 0.02 | 0.005 | 26.299 |

**Supplementary Table S6.** Held-out cohort replication. The recoverability measurements repeated on two single-cell cohorts that contributed nothing to the analysis set: a 231-patient anti-PD-1-treated non-small cell lung cancer atlas (NSCLC; Liu et al., GEO accession GSE243013) and a 15-patient head and neck squamous cell carcinoma cohort treated with anti-PD-L1 and anti-CTLA-4 (HNSCC; Cha et al., GEO accession GSE286827). Each cohort was normalized and deduplicated exactly as the analysis set, every rearrangement also present in the canonical set was removed, and 10,000 of the remainder were sampled per cohort-chain at a fixed seed (three cells: NSCLC $\alpha$, NSCLC $\beta$ and HNSCC $\beta$, the head and neck cohort contributing $\beta$ only); sequences whose OLGA $P_{\text{gen}}$ was zero were excluded, which is why the scored count falls below 10,000 for NSCLC $\alpha$ (221 dropped) and NSCLC $\beta$ (1). The last row is the adjusted Rand index between the confusion grouping derived independently within the cohort and the grouping derived from the analysis set, over the genes shared by both; for comparison, the analysis-set grouping agrees with the IMGT family partition at 0.0519 ($\alpha$) and 0.214 ($\beta$).

| | $\alpha$ | | $\beta$ | | |
|---|---:|---:|---:|---:|---:|
| **quantity** | **analysis set** | **NSCLC** | **analysis set** | **NSCLC** | **HNSCC** |
| sequences scored | 37,687 | 9,779 | 80,409 | 9,999 | 10,000 |
| V-gene conditional entropy (nats) | 1.2925 | 1.2851 | 2.7055 | 2.4365 | 2.4565 |
| J-gene conditional entropy (nats) | 0.0115 | 0.0080 | 0.0474 | 0.0267 | 0.0256 |
| top-1 V coverage | 0.493 | 0.483 | 0.203 | 0.328 | 0.332 |
| top-10 V coverage | 0.847 | 0.872 | 0.607 | 0.727 | 0.729 |
| top-20 V coverage | 0.921 | 0.935 | 0.881 | 0.899 | 0.888 |
| top-1 mass $\geq$ 0.5 | 0.570 | 0.574 | 0.134 | 0.230 | 0.224 |
| top-1 mass $\geq$ 0.99 | 0.132 | 0.128 | 0.042 | 0.080 | 0.080 |
| V genes with $\geq$ 20 sequences | 44 | 41 | 47 | 43 | 43 |
| ARI vs analysis-set grouping | — | 0.959 | — | 0.867 | 0.815 |
